# Use of Integrative Proteomic Analysis to Uncover Shared Molecular Signatures Across Distinct Forms of Hydrocephalus

**DOI:** 10.64898/2026.11.05.764703

**Authors:** Gowthami Mahendran, Piriyankan Kirupaharan, David Audu, Bonnie L. Blazer-Yost

## Abstract

Hydrocephalus is a heterogeneous disease characterized by disruption of cerebrospinal fluid (CSF) dynamics and can arise from diverse etiologies, including post-hemorrhagic hydrocephalus (PHH), hydrocephalus following subarachnoid hemorrhage (SAH), post-infectious hydrocephalus (PIH), and idiopathic normal pressure hydrocephalus (iNPH). Hydrocephalus has high disease burden in pediatric and older populations and a marked increase in prevalence with advancing age. Despite this high incidence, the underpinning molecular and signaling pathway alterations remain incompletely understood. Hence CSF proteomic profiling would enable a robust and high-resolution approach for exploring disease-specific biomarkers and to discover shared molecular mechanisms underlying hydrocephalus. In this study, we performed a cross-study proteomic analysis to evaluate biomarker proteins reported across PHH, SAH, PIH, and iNPH groups. Using published proteomic datasets, we comparatively analyzed CSF proteins to investigate the differentially expressed proteins (DEPs), their common molecular signatures, and enriched biological pathways across hydrocephalus groups. Across etiologies, recurrently altered proteins were predominantly associated with inflammatory signaling (cytokine, chemokine, interleukin-(IL-17), tumor necrosis factor (TNF)), toll-like receptor (TLR) signaling, blood-brain-barrier (BBB) disruption leading to coagulation cascades, and neuronal injury responses. Hemorrhage-associated forms of hydrocephalus (PHH and SAH) demonstrated enriched coagulation and hemoglobin-related pathways, whereas infection associated forms (PIH) showed enhanced immune and inflammatory responses. iNPH exhibited alterations in proteins linked to neurodegeneration, synaptic function, and impaired CSF clearance. Interestingly, our cross-study analyses identified recurrently dysregulated proteins across hydrocephalus subtypes. Collectively, our cross-study proteomic analyses define shared and etiology-specific proteomic signatures across hydrocephalus subtypes and identify potential CSF-based biomarkers for diagnostic stratification and future precision therapeutic approaches.

## 1. Introduction

Hydrocephalus is a severe neurological condition categorized by abnormal accumulation of cerebrospinal fluid (CSF) within the brain ventricles, resulting in ventricular enlargement. Hydrocephalus, arises from diverse etiologies leading to variable clinical outcomes **[26, 29]**. The choroid plexus epithelial cells (CPe), located in the lateral, third and fourth ventricles, are responsible for producing around 80% of CSF, with the remaining 20% derived from interstitial fluids regulated by the blood-brain barrier (BBB) **[13]**. Produced CSF then flows through ventricles, subarachnoid space and parenchyma before being absorbed into venous circulation **[13, 49, 58]**. In a healthy human, normal CSF production is approximately 0.5 liters per day, with the total CSF volume being renewed three to four times daily **[12, 13]**. Disruption of the tight regulation of CSF can contribute to the development of hydrocephalus which involves changes to the CSF dynamics, presented with either increased or normal intracranial pressure (ICP) **[25]**. When CSF flow is disrupted by a physical obstruction within the ventricular system, commonly at the cerebral aqueduct or ventricular foramina, the condition is classified as noncommunicating (obstructive) hydrocephalus, which results in the dilation of the upstream ventricles **[41]**. In contrast, communicating hydrocephalus develops when ventricular pathways remain clear but CSF accumulation can result from impaired clearance or overproduction of CSF **[5, 31]**. Idiopathic normal pressure hydrocephalus (iNPH) is a communicating type of hydrocephalus characterized by ventricular enlargement despite normal or intermittently elevated ICP. Regardless of classification, hydrocephalus ultimately reflects an imbalance between CSF production, circulation, and absorption within the ventricular and subarachnoid spaces **[23, 28, 42]**.

In addition to the mechanistic classification, hydrocephalus can also be categorized by origin as primary (genetic, congenital, or developmental) or acquired, the latter arising from conditions such as subarachnoid hemorrhage (SAH) or intraventricular hemorrhage (IVH), infection, trauma, or tumor **[50]**. Both primary and acquired forms may manifest as non-communicating or communicating hydrocephalus. However, these conventional classifications do not fully capture the complex and evolving pathophysiology of hydrocephalus. More recently, a study published the pulsatile nature in CSF movement across the ventricles that was explained through cerebral windkessel (CWK) mechanism, shifting from the traditional way of classifying hydrocephalus as obstruction to the subarachnoid space. This CWK model offers an alternative framework for intracranial fluid dynamics, suggesting that hydrocephalus may arise from impaired transmission of pulsatile energy within CSF pathways rather than primarily through impaired CSF absorption **[16]**. Recent advances have expanded the traditional view of hydrocephalus as an imbalance between CSF production and absorption toward a more integrated framework that considers altered CSF pulsatility, impaired intracranial compliance, and disrupted fluid dynamics. Emerging evidence further implicates glymphatic and perivascular clearance dysfunction, cerebrovascular abnormalities, BBB and blood-CSF barrier (BCSFB) disruption, and neuroinflammation in hydrocephalus pathophysiology **[4, 21, 72]**. Altered glymphatic function may impair CSF–interstitial fluid exchange and waste clearance, while the central nervous system (CNS) barrier disruption can alter molecular exchange between the blood, brain, and CSF compartments. These changes along with sustained inflammatory signaling, influence choroid plexus function and CSF homeostasis, highlighting the complex and interconnected mechanisms underlying hydrocephalus. These diverse yet intersecting mechanisms emphasize the complex and heterogeneous nature of hydrocephalus, highlighting the necessity of a deep understanding of the disease mechanism for therapeutic targeting.

Despite its significant clinical impact, there are currently no long-term, effective pharmaceutical treatments available to treat hydrocephalus. Management primarily relies on surgical interventions, such as CSF shunting, endoscopic third ventriculostomy (ETV), or ETV with choroid plexus cauterization which are associated with complications including infection, obstruction, shunt failures and the need for frequent revisions **[45, 60, 65, 68]**. Consequently, there remains a substantial unmet need for safer, more durable and noninvasive therapeutic strategies. Hence, the continued dependence on surgical methods to treat hydrocephalus highlights the critical gap in knowledge of CSF regulation, and the biological processes and pathways linked to CSF production and clearance. The development of robust biomarkers for hydrocephalus is critical for supporting early detection, longitudinal monitoring, and individualized treatment strategies, particularly in predicting shunt outcomes. Approaches that enable comprehensive profiling of disease-associated changes at the protein level offer a promising avenue to uncover coordinated signaling events that may drive pathogenesis and inform targeted intervention strategies.

CSF, a biological fluid rich in many proteins (cytokines, inflammatory mediators, growth factors, enzymes), metabolites, and lipids. It closely reflects the CNS physiology and acts as an ideal source of biomarker for hydrocephalus due to its direct interaction with the ventricular system, and choroid plexus **[30, 71]**. Additionally, an added advantage of CSF profiling is that CSF can be extracted from living patients which can provide a clinically accessible window of investigating the disease-associated molecular alterations associated with CSF dysregulation. Current advances in high-throughput technologies now facilitate unbiased CSF proteomic profiling, allowing the detection of functional protein-level alterations. However, despite multiple proteomic investigations across hydrocephalus types, to the best of our knowledge, no study has compared and reported which proteins and molecular pathways are recurrently dysregulated across PHH, SAH-associated hydrocephalus, PIH, iNPH, and other hydrocephalus subtypes, and whether these shared signatures could serve as biomarkers or reveal common disease mechanisms.

Therefore, in this work, we have systematically examined the published and publicly available original proteomic studies over the past 16 years, characterizing the proteomes of different hydrocephalus subtypes to identify protein biomarkers associated with various forms of hydrocephalus. Our objective was to cross-analyze reported proteomic signatures across independent cohorts and hydrocephalus subtypes to identify recurrently dysregulated proteins and shared molecular patterns that may reflect core disease mechanisms and serve as candidate biomarkers or therapeutic targets. The proteomics studies selected and analyzed in this study were conducted in accordance with the Preferred Reporting Items for Systematic Reviews and Meta-Analyses (PRISMA) guidelines **[46]** to ensure methodological rigor, transparency, and reproducibility.

## 2. Materials and Methods

### 2.1. Data collection

The aim of our study was to gain a broad understanding of how proteomics in hydrocephalus has been studied across different hydrocephalus subtypes to find common biomarker proteins. Three prominent databases were chosen (PubMed, Scopus, and Web of Science), covering the period from 2010 to 2026 and the keywords for searches used were “Hydrocephalus” AND “Proteomics”. Search results from these databases were imported into Covidence, which was used to manage references, remove duplicates, and to facilitate the screening process. Two reviewers independently screened titles and abstracts, followed by full-text assessment based on predefined inclusion and exclusion criteria described below. Discrepancies were resolved through consensus. A total of 20 studies met the inclusion criteria and were included in the final analysis.

### 2.2. Eligibility criteria and study selection for data analysis

Our initial search yielded 172 papers: PubMed (n = 81), Scopus (n = 46), and Web of Science (n =45). 75 duplicate papers were removed, resulting in 97 papers eligible for the initial screening phase (i.e., title and abstract screening). After retrieving the papers, we included those that met the following criteria. Our inclusion criteria were, papers written in English and peer reviewed, papers that focus on different types of hydrocephalus and proteomics, papers that performed proteomics investigations, papers published in the last 16 years, papers that compared the hydrocephalus subtypes with their relevant age-matched healthy controls. Our exclusion criteria were, papers not written in English or non-peer reviewed (preprints or bio archives), papers that focus on hydrocephalus but not proteomics and vice versa, papers that are not research articles (reviews, opinions, editorials etc.), papers that are only focusing on the proteomics of the treatment responses (e.g., shunt responsiveness vs controls), papers that compared the hydrocephalus subtypes with other neurological conditions (e.g., iNPH vs Alzheimer’s diseases (AD), iNPH vs mild cognitive impairment (MCI), traumatic brain injury (TBI) vs iNPH).

From these 97 papers, 57 papers were excluded based on our exclusion criteria in the initial title and abstract screening. This left 40 papers for second phase of review. 20 papers were excluded further because of the wrong intervention, or study design (compared the shunt responsiveness), or patient population (not compared against the healthy controls) or their major focus was not hydrocephalus (primary focus was on a different neurological condition). A final set of 20 papers met all inclusion criteria following the full-text screening and were included in this study (Table A1). Figure 1 presents the PRISMA flow diagram of the screening process for this study analysis.

**Table A1.** Hydrocephalus proteomics studies from literature included in this study.

| **Study Model** | **Hydrocephalus Type** | **Proteomics Method Used** | **Published Year** | **Ref** |
|---|---|---|---|---|
| Human CSF (vCSF) | Congenital, tumor related hydrocephalus | Microcapillary LC-MS/MS | 2010 | [67] |
| Human CSF (vCSF) | PHH | MS (LFQ) | 2012 | [43] |
| Human CSF (LP CSF) | iNPH | HPLC coupled to LC-TOP-MS | 2017 | [44] |
| Human (brain biopsies) | TBI/ PHH | MS (LFQ/ Stable isotope dimethyl labeling) | 2018 | [1] |
| Human CSF (vCSF) | PHH (SAH) | MALDI-TOF | 2019 | [59] |
| Rat CSF (vCSF) | Congenital Hydrocephalus | LC-MS/MS | 2021 | [51] |
| Human CSF (LP CSF) | iNPH | LC-MS/MS | 2021 | [61] |
| Human CSF (SAH, trauma, tumors) (not mentioned) | Communicating hydrocephalus (non-obstructive) | LC-MS/MS | 2021 | [70] |
| Human CSF (LP CSF) | SAH (PHH) | MS | 2022 | [17] |
| Human CSF (vCSF) | SAH (PHH) | PEA on O-link | 2022 | [37] |
| Human CSF (LP CSF) | iNPH | PEA on O-link | 2023 | [7] |
| Human CSF (LP CSF) | PIH | MS (LFQ) | 2023 | [9] |
| Rats (Lateral ventricle ChP) | PIH and PHH | LC-MS/MS (TMT) | 2023 | [53] |
| Human CSF (LP CSF) | iNPH | LC-MS/MS | 2023 | [55] |
| Human CSF (LP CSF) | iNPH shunt responsive | PEA on O-link | 2024 | [30] |
| Human serum and urine | PHH (IVH) preterm neonates | PEA on O-link | 2024 | [64] |
| Human CSF (vCSF) | PHH (SAH) | LC-MS/MS | 2024 | [38] |
| Human CSF (LP CSF) | PHH, NMDARE, pediatric brain-tumor associated hydrocephalus | PEA on O-link | 2025 | [8] |
| Human CSF (LP CSF) | PHH | MS (LFQ) | 2025 | [10] |
| Human CSF (LP CSF) | iNPH | MS (DIA) | 2025 | [22] |
| Mouse CSF (vCSF) | PHH (LPA injected) | LC-MS/MS (TMT) | 2025 | [57] |
| Human CSF (LP CSF and vCSF) | iNPH | LC-MS/MS | 2025 | [69] |

NMDARE: Anti-N-methyl-D-aspartate receptor, LPA: Lysophosphatidic acid, LFQ: label-free quantitation, PEA: proximity extension assay, LC-MS/MS: Liquid chromatography-tandem mass spectrometry, HPLC: High-performance liquid chromatography, LC-TOP-MS: Top-down Liquid Chromatography-Mass Spectrometry, TMT: tandem mass tag, DIA: Data-independent acquisition, MALDI: Matrix-Assisted Laser Desorption/Ionization.

**Figure 1.**
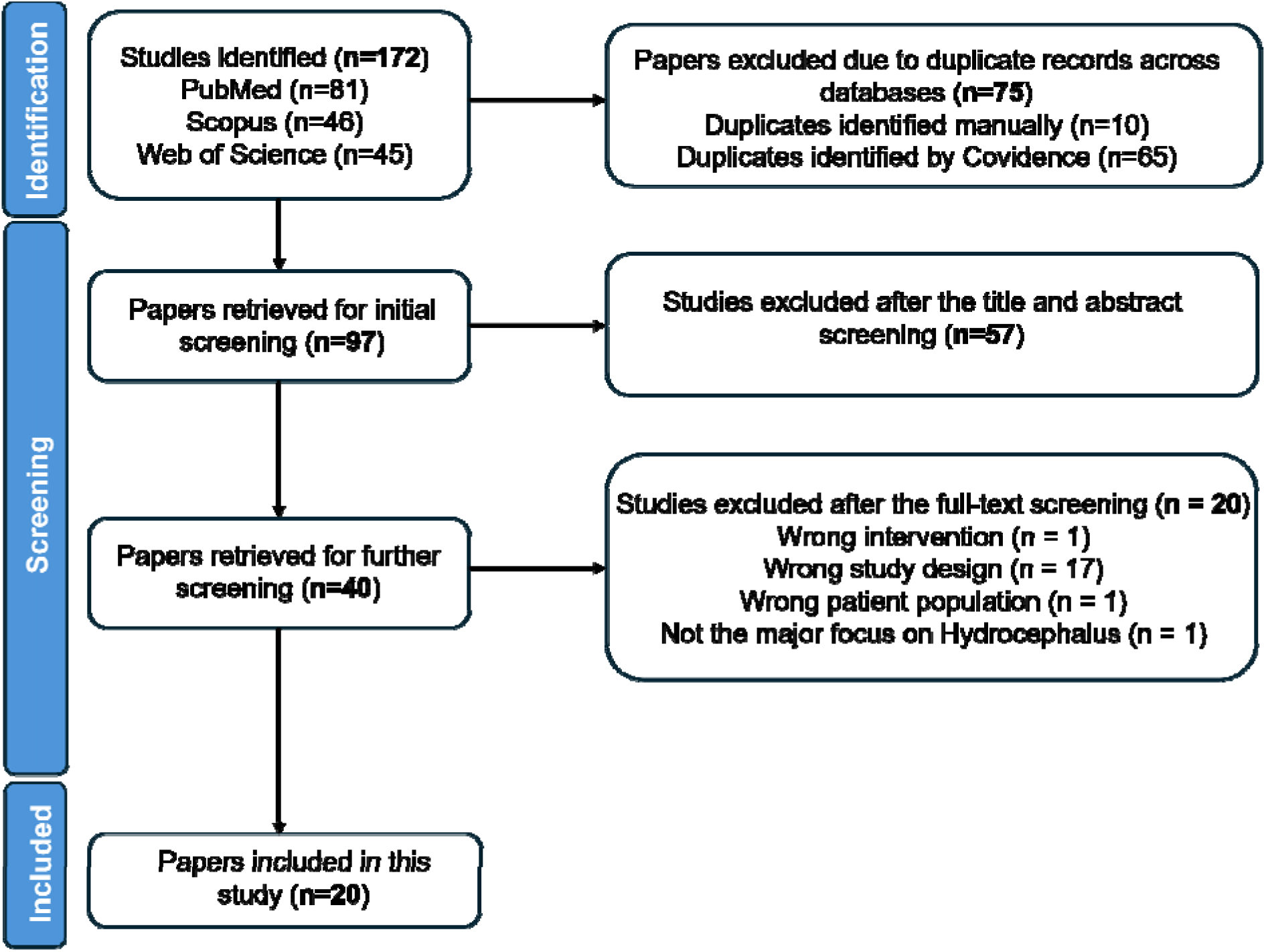
PRISMA 2020 flow diagram summarizing the identification, screening (based on the eligibility criteria) and inclusion of final numbers of the study. The number of studies included at each step are shown within the brackets. A flow diagram was developed with Covidence assistance.

### 2.3. Differentially expressed protein (DEP) identification and pathway analysis

From the finalized 20 studies, we extracted the differentially expressed proteins (DEPs) from papers that had published processed proteomics data file which were publicly available or received upon request from the author. We selected the DEPs based on these two cutoffs: 0.5 ≤ log_2_ Fold change ≤ -0.5 and p value ≤ 0.05. Upregulated and downregulated proteins identified based upon this predefined cutoff are listed in Table S1. The top five up and downregulated candidates from each subtypes are given in Table A2 briefly. The DEPs belonging to the four subtypes highlighted in this study (PHH vs control, SAH vs control, iNPH vs control and HBM vs control) are given in separate tabs. Note that HBM refers to bacterial meningitis-associated hydrocephalus, which falls within the PIH subtype. Because only one PIH proteomic study was identified and it specifically investigated HBM, this category is referred to as HBM rather than PIH throughout the analysis. Further, a Venn Diagram was created separately for upregulated and downregulated proteins for the four hydrocephalus subtypes focused on this study to investigate the overlapping and unique proteins for each subtype. The overlapping proteins identified through the Venn diagram analysis are listed in Table S2. R studio was utilized to generate the Venn diagram across the four groups. ShinyGO 0.85 online tool was used to produce pathway maps for the gene ontology (GO) terms: biological pathways (BP), molecular function (MF) and cellular components (CC) and Kyoto Encyclopedia of Genes and Genomes (KEGG) pathways **[20]**. The total DEPs for each subtype were individually uploaded into ShinyGO tool to generate the relevant pathway plots. For relevance, we considered only the top ten signaling pathways associated with these proteins, ranked according to false discovery rate (FDR) and fold enrichment score. For each subtype, four enrichment plots were generated: biological process (BP), molecular function (MF), cellular component (CC), and KEGG pathways. The output files of associated proteins to these pathways from ShinyGO are included in Table S3.

**Table A2.** Top five up and downregulated proteins from each hydrocephalus subtypes with their respective fold change in expressions.

| **Gene Symbol** | **Full Protein Name** | **Protein Function** | **Fold Change** |
|---|---|---|---|
| **PHH vs Control** |  |  |  |
| NELL2 | Neural epidermal growth factor-like protein 2 | Signaling molecule in neuronal development | -2.67 |
| AGRN | Agrin | An ECM protein mediating acetylcholine receptor clustering at the neuromuscular junction | -1.75 |
| KLK7 | Kallikrein-related peptidase 7 | A role in degrading amyloid-β plaque | -1.61 |
| BANK1 | B-cell scaffold protein with ankyrin repeats 1 | A scaffold protein negatively regulating CD-40 mediated Akt signaling | -1.37 |
| IL20RA | Interleukin-20 receptor subunit alpha | Signaling involved in inflammatory responses | -1.11 |
| CHI3L1 | Chitinase-3-like protein 1 | Key inflammatory protein secreted by microglia and astrocytes | 10.05 |
| FTL | Ferritin light chain | Iron storage protein, builds up in the brain | 10.01 |
| CCL22 | C-C motif chemokine 22 | Cytokine secreted by macrophages, dendritic cells and B- cells | 9.47 |
| FABP1 | Fatty acid-binding protein, liver | Impacting neuroinflammatory processes | 9.13 |
| COL9A1 | Collagen alpha-1(IX) chain | Extracellular matrix structural proteins | 8.85 |
| **SAH vs Control** |  |  |  |
| CHGB | Chromogranin B | Key regulator of calcium in the brain | -2.30 |
| PTGDS | Prostaglandin D synthase, lipocalin-type | Neuromodulator and a neuroprotectant | -2.03 |
| THY1 | Thy-1 cell surface glycoprotein | Highly expressed in mature neurons | -2.00 |
| APLP1 | Amyloid beta precursor–like protein 1 | Synaptic adhesion molecule | -1.93 |
| SCG3 | Secretogranin III | A role in neurotransmitter release | -1.88 |
| LIF | Leukemia Inhibitory Factor | Anti-inflammatory and a neuroprotective cytokine | 5.30 |
| MCP-3 | Monocyte Chemoattractant Protein-3 | Inflammatory chemokine driving inflammatory pathway after an injury | 3.40 |
| TTR | Transthyretin | Synthesized in choroid plexus and secreted into CSF | 3.20 |
| IL-6 | Interleukin-6 | Acts as both pro- and anti-inflammatory cytokine | 3.10 |
| OSM | Oncostatin M | A role in neuroinflammatory pathways, and increases the production of IL-6 and MCP-1 | 2.60 |
| **iNPH vs Control** |  |  |  |
| MSLNL | Mesothelin-like protein | Pseudogene | -2.50 |
| CCND2 | G1/S-specific cyclin-D2 | Cell regulator essential for cell proliferation and neurogenesis | -2.48 |
| SLC1A4 | Neutral amino acid transporter A | Transporting L-serine across the blood-brain barrier and from astrocytes to neurons | -2.31 |
| MYOM2 | Myomesin-2 | Embryonic brain development | -2.27 |
| CBLN1 | Cerebellin-1 | Maintaining excitatory synapses | -2.25 |
| GBP2 | Guanylate-binding protein 2 | High expression in microglia and linked to neuroinflammation | 3.59 |
| PMS1 | PMS1 protein homolog 1 | Linked to different neurological diseases | 2.71 |
| FAT3 | Protocadherin Fat 3 | Essential for brain and synaptic development | 2.62 |
| CLPX | ATP-dependent clpX-like chaperone, mitochondrial | Maintains mitochondrial health and linked to neuro diseases | 2.39 |
| VIM | Vimentin | A key marker of neuroinflammation | 2.36 |
| **HBM vs Control** |  |  |  |
| TNFRSF13 | Tumor necrosis factor receptor superfamily member 13B | Onco-inflammatory marker in the brain | 14.43 |
| S100A8 | S100 calcium-binding protein A8 | A key calcium-binding protein that functions as a damage-associated molecular pattern (DAMP), promoting neuroinflammation in conditions such as TBI | 11.16 |
| UBE2D1 | Ubiquitin-conjugating enzyme E2 D1 | A key ubiquitin-conjugating enzyme in the brain | 10.99 |
| AZU1 | Azurocidin 1 | Inflammatory modulator with high expression in neurons | 10.83 |
| C4BPB | Complement component 4 binding protein beta chain | Biomarker for neuroinflammatory diseases | 10.13 |

## 3. Results

### 3.1. Proteomic investigation of hydrocephalus has expanded with human CSF studies concentrated in PHH, SAH and iNPH

Following PRISMA-based study selection, a total of 20 proteomic studies were included in our analysis, and their distribution across hydrocephalus subtypes, publication year, sample type, and model system are summarized in Figure 2. The heatmap demonstrates that proteomic investigations in hydrocephalus have increased notably in recent years, with most studies published after 2020 (Figure 2A). PHH/IVH, SAH and iNPH were the most frequently represented hydrocephalus subtypes across the analyzed proteomic studies within the last 16 years. Among these hydrocephalus subtypes, PHH, including cases associated with SAH and IVH, represents the most frequently studied category, with a marked rise in studies between 2021 and 2025. Similarly, iNPH has also gained increasing attention in recent years, reflecting growing interest in adult and communicating forms of disease. Figure 2B highlights the predominance of CSF (both lumbar puncture CSF (LP CSF) and ventricular CSF (vCSF)) as the primary biological sample used for proteomic profiling, accounting for many included studies. In contrast, serum, choroid plexus tissue (ChP), and urine were evaluated in only a limited number of investigations. Interestingly, most studies on CSF proteomics were from human samples, with minimal utilization of animal models such as mice or rats. Collectively, these findings indicate that hydrocephalus proteomic research over the past 16 years has been largely centered on human CSF-based biomarker discovery, with increasing focus on hemorrhage-associated and iNPH subtypes, whereas congenital, infectious hydrocephalus studies remain relatively limited.

**Figure 2.**
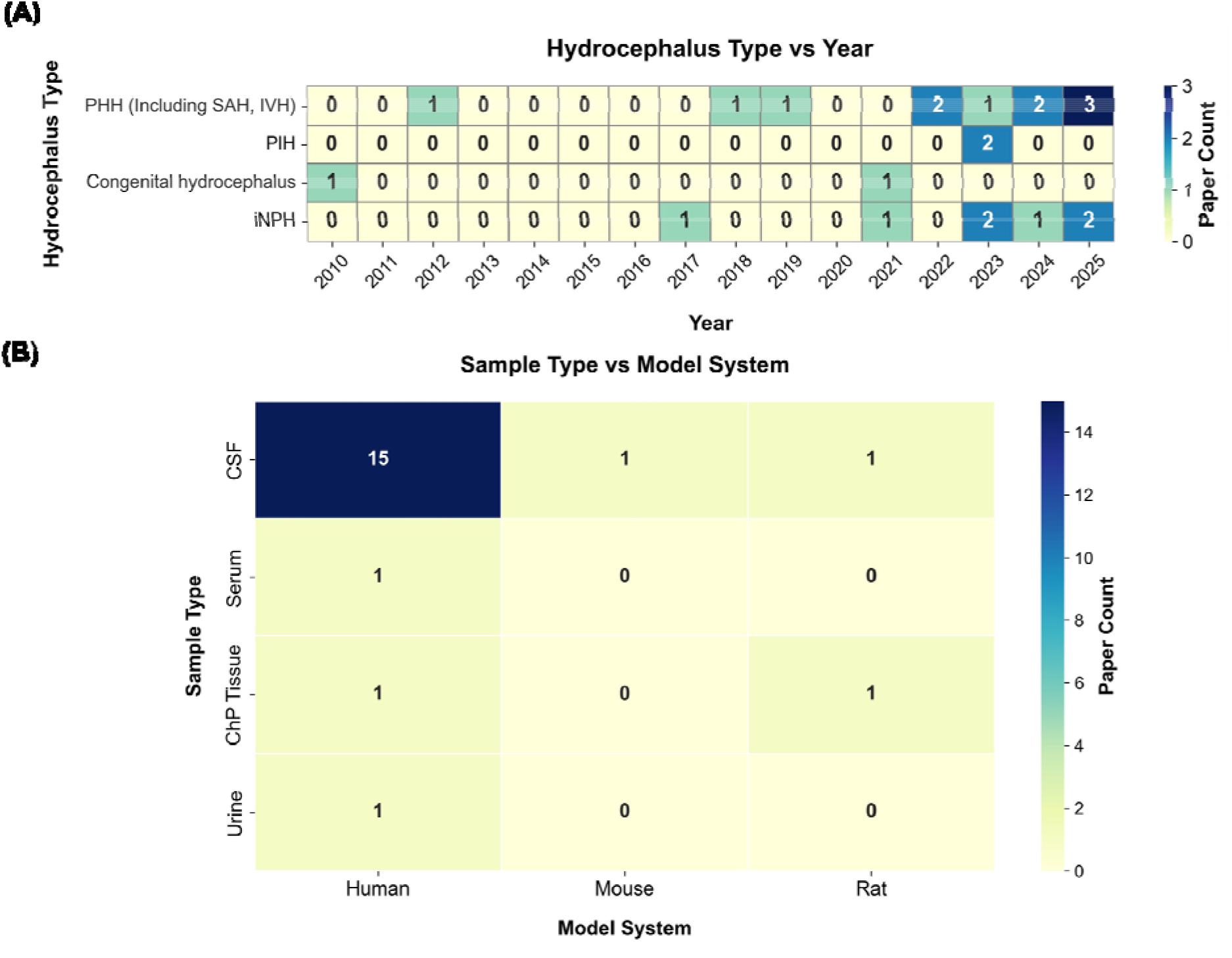
Heatmap illustrations of (A) proteomics studies across different hydrocephalus types over the past 16 years (2010 to 2026) and (B) sample vs model system plot showing the studies conducted in different model systems (human, rat and mouse) using different sample types (CSF, serum, ChP tissue and urine).

We investigated subtype-specific protein signatures and identified both unique and shared protein candidates across different hydrocephalus subtypes. Unless otherwise specified, all biomarkers discussed in this study were identified through comparisons with healthy controls from published datasets that could possibly serve as biomarkers for each hydrocephalus category. Further, in our study, we focused on two different CSF samplings (LP CSF and vCSF), and two different proteomics technique (mass spectrometry, and proximity extension assay (PEA) with O-link). By comparing the findings from independent studies, our goal is to discover recurrent molecular patterns that may represent common disease mechanisms and subtype-specific biological changes. Further, we analyzed the functional networks associated with these proteins to better understand their roles in CSF dysregulation, neuroinflammation, ciliary dysfunction, barrier integrity, and other related pathophysiological processes.

### 3.2. Most common DEPs are linked to coagulation cascades, inflammatory responses, immune responses and cytokine signaling in different forms of hydrocephalus

PHH can arise following various intracranial hemorrhagic events and is often a serious complication in premature infants (born before 32 weeks of gestation) **[15]**. In the preterm population, PHH most commonly develops secondary to germinal matrix hemorrhage (GMH), IVH, SAH and subsequent post-hemorrhagic ventricular dilation (PHVD) **[47]**. In adults, PHH commonly develops following intracranial hemorrhage, particularly SAH and IVH, including hemorrhage associated with traumatic brain injury (TBI) **[11]**. These conditions can lead to long-term neurological sequelae, including cognitive delay, visual impairment, motor dysfunction, cerebral palsy, and behavioral dysfunction **[36]**. The PHH studies identified in our search primarily involved preterm infants with GMH or IVH, whereas the SAH-associated hydrocephalus studies predominantly involved adult patient populations. Therefore, to ensure clear and biologically appropriate classification, PHH and SAH-associated hydrocephalus were analyzed separately. Further, to characterize the functional relevance of the candidate biomarker proteins reported in PHH from the literature, we performed pathway enrichment analysis. The GO and KEGG pathway enrichment plots highlight the overrepresented signaling pathways associated with DEPs we included from our study pool, with the higher -log_10_(false discovery rate or FDR) and fold enrichment indicating the most statistical significance and increased abundance, respectively. KEGG pathway results for PHH demonstrated the strongest enrichment in pathways related to inflammatory signaling, extra cellular matrix (ECM) regulation, cellular communication, and injury response. Several interleukins, mitogen-activated protein kinases (MAPKs), and vascular endothelial growth factors (VEGFs) were mapped to key signaling cascades, including the Janus Kinase–Signal Transducer and Activator of Transcription (JAK–STAT), MAPK, and Phosphoinositide 3-kinase–Protein Kinase B (PI3K–Akt) pathways which were included in the top 10 pathways (Figure 3A, left). Similarly, the GO BP enrichment analysis revealed that the top enriched pathways were related to inflammatory and cytokine-mediated responses, including multiple cytokines and chemokines (Figure 3A, right). These GO enrichment findings are consistent with the KEGG pathway analysis and support the identified roles of the DEPs in PHH/IVH, highlighting a strong inflammatory and immune-related component underlying the condition.

**Figure 3.**
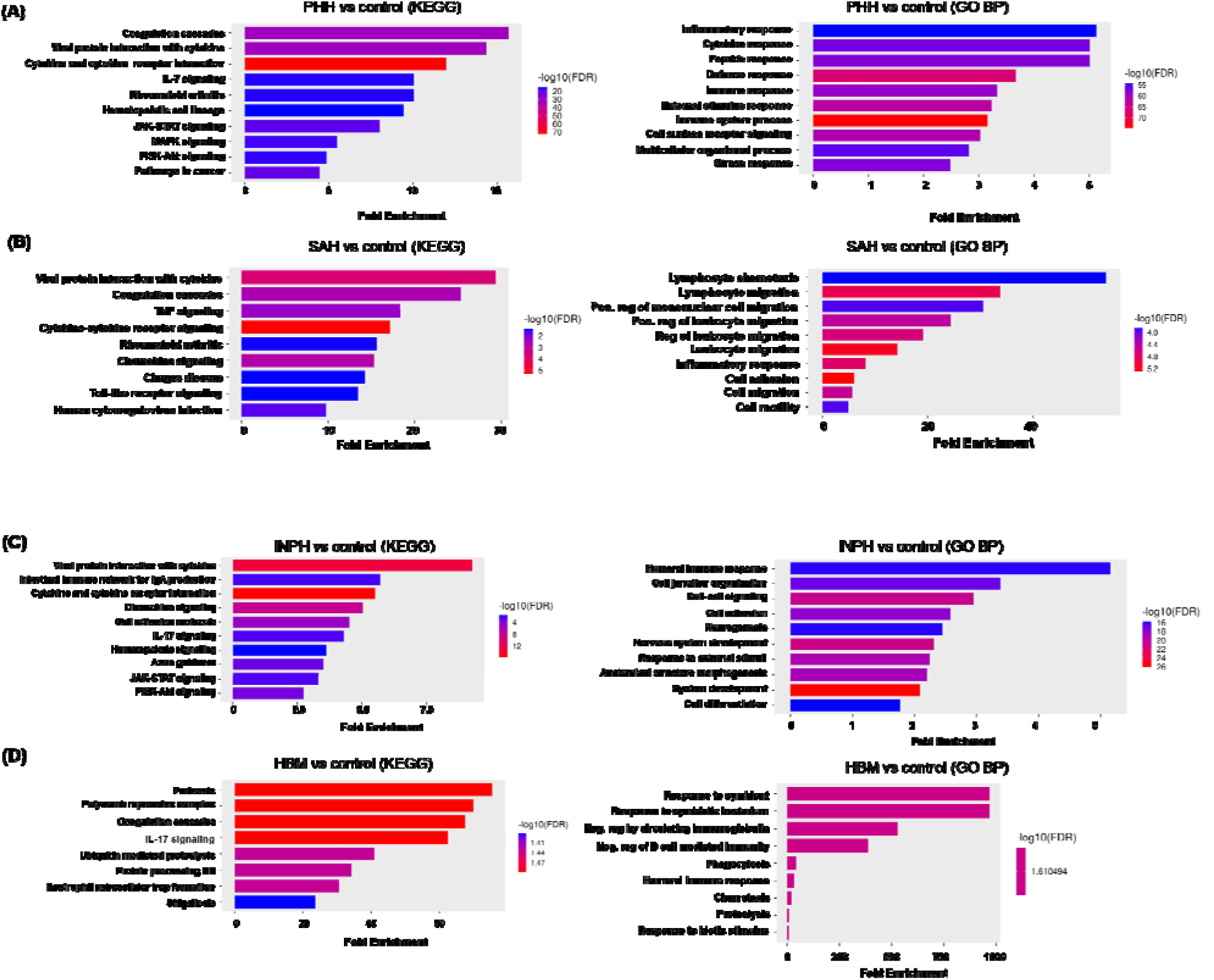
Top ten highly enriched KEGG (left) and GO term (right) pathways of (A) PHH, (B) SAH, (C) iNPH and (D) HBM. Pathway diagrams were generated using Shiny GO online tool.

Beyond IVH-associated PHH in the developing brain, hydrocephalus can also develop following SAH, a severe form of hemorrhagic stroke associated with substantial morbidity and mortality **[34]**. A major pathological consequence of SAH is the accumulation of blood and its secondary breakdown products, including coagulation factors and degradation products, within the CSF, which contributes to secondary brain injury **[3]**. Blood coagulation within the subarachnoid space causes dysregulation in CSF dynamics **[3]**. Consistent with the close interplay between coagulation and inflammation in SAH pathology, our KEGG pathway analysis demonstrated strong enrichment of cytokine, tumor necrosis factor (TNF), chemokine, Toll-like receptor (TLR), and coagulation signaling pathways, indicating that SAH triggers coordinated activation of inflammatory and coagulation-related processes, thereby highlighting the central role of immune–coagulative crosstalk in driving secondary brain injury in SAH patients (Figure 3B, left). In contrast, the GO BP analysis revealed enrichment of pathways related to cellular migration, adhesion, and motility in the SAH condition (Figure 3B, right). This enrichment suggests increased leukocyte recruitment, endothelial activation, and cytoskeletal remodeling in response to blood exposure in the subarachnoid space. Although the two analyses highlighted distinct pathway categories, these findings are complementary as KEGG primarily represents established molecular signaling pathways, whereas GO BP demonstrates broader biological and cellular processes. Thus, these processes are consistent with neuroinflammation and cytokine signaling, both of which contribute to secondary brain injury following SAH.

The third subtype of hydrocephalus is iNPH, another form of hydrocephalus that affects adults, is a progressive but a reversible neurological disorder characterized by ventriculomegaly, gait abnormality, urinary incontinence, dementia and cognitive defects **[2, 25]**. It is often misdiagnosed because of its overlapping clinical symptoms with other neurodegenerative conditions, such as Alzheimer’s disease (AD), Parkinson’s disease (PD), and mild cognitive impairment (MCI), making the diagnosis highly challenging **[52]**. Established CSF biomarkers, including amyloid-β42 and tau proteins, are widely used in the evaluation of AD and overlapping cognitive symptoms in iNPH **[33]**. However, these biomarkers lack specificity for iNPH due to shared pathological features with AD. Further to gain insight into the biological pathways of these proposed biomarkers in iNPH, pathway enrichment analysis was performed. KEGG pathway results exhibited the strongest enrichment in pathways related to inflammatory signaling (cytokine, chemokine signaling, IL-17 signaling). In addition, pathways related to cell adhesion, axon guidance, and intracellular signaling cascades such as JAK– STAT and PI3K–Akt were prominently enriched. The enriched inflammatory and adhesion-related pathways support the biological relevance of the identified biomarkers and aligns with the established pathological features of iNPH, including neuroinflammation, ependymal dysfunction, and impaired synaptic and axonal integrity (Figure 3C, left). Similarly, the GO BP enrichment analysis revealed that the top enriched pathways were related to immune response, cell signaling, cell adhesion and neuronal system development (Figure 3C, right). Overall, the KEGG and GO enrichment results for iNPH reflect an integrated network of inflammatory activation and impaired cell and axonal signaling contributing to iNPH progression.

Lastly, PIH arises as a consequence of CNS infections that trigger intense inflammatory responses within the meninges and ventricular system, ultimately disrupting CSF circulation and absorption **[32, 66]**. Infectious etiologies such as tuberculous meningitis, neurocysticercosis, and bacterial or viral meningitis promote the accumulation of inflammatory responses, ependymal injury, arachnoid fibrosis, and obstruction of CSF pathways, leading to either communicating or obstructive hydrocephalus **[48, 62]**. In infants, PIH commonly develops secondary to antenatal infections (e.g., TORCH infections) or postnatal infectious events such as sepsis and meningitis, both of which are associated with ventriculitis and long-term impairment of CSF resorption mechanisms **[19]**. Emerging evidence further demonstrates that PIH and PHH exhibit overlapping inflammatory cytokine signatures, including elevations in pro-inflammatory mediators and chemokines, suggesting shared immunopathological mechanisms driving ventricular enlargement **[24, 35, 53]**. Despite these mechanistic insights, standardized diagnostic criteria for PIH in adults remain lacking. Thus, to further investigate the biological significance of these inflammatory protein alterations, we performed our pathway analysis based on the HBM proteomics dataset. The KEGG enrichment revealed significant overrepresentation of pathways linked to pertussis, shigellosis, and cytokine signaling (notably IL-17 signaling) in HBM patients. Rather than indicating pathogen-specific infection, these enrichments highlight shared innate immune and pro-inflammatory signaling pathways as prominent molecular features of HBM-associated hydrocephalus (Figure 3D, left). To further support the KEGG results, GO BP enrichment analysis demonstrated strong activation of host immune response pathways in HBM, particularly those related to response to symbiont, response to biotic stimulus, and phagocytosis (Figure 3D, right). Enrichment of humoral immune response and immunoglobulin-mediated regulation further indicates strong involvement of both innate and adaptive immunity within the CSF. Together these findings underscore immune-driven inflammation and host–pathogen interactions as central molecular mechanisms underlying HBM-associated hydrocephalus (Figure 3D, left).

### 3.3. Cross-subtype comparison of recurrent proteins reveal DEPs and pathways linked to cytokine responses

The prime goal of our study is to identify shared proteomic signatures across selected hydrocephalus subtypes and to characterize the biological and signaling pathways associated with these overlapping proteins. To systematically evaluate cross-subtype similarity, a Venn diagram was created to determine shared upregulated and downregulated protein sets. As illustrated in Figure 4A, 39 upregulated proteins were common between iNPH and PHH, whereas only five proteins overlapped between PHH and SAH. Notably, a single protein (CCL4) was shared across all three subtypes (iNPH, PHH, SAH), and no overlap was observed with the HBM group. In contrast, analysis of the downregulated protein set (Figure 4B) revealed more modest overlap, with three proteins shared between iNPH and PHH and six shared between iNPH and SAH (Table A3). Importantly, no protein was consistently downregulated across all subtypes. This observation was further substantiated by the heatmap in Figure 4C, which clearly illustrates the overlapping proteins across subtypes, with fold-change intensity depicted by the color scale.

**Table A3.** Upregulated and downregulated DEPs from overlapping hydrocephalus subtypes.

| **Gene Symbol** | **Full Protein Name** | **Up or Down** |
|---|---|---|
| **iNPH vs SAH** |  |  |
| CHGA | Chromogranin-A | Down |
| CX3CL1 | C-X3-C motif chemokine ligand 1 | Down |
| NRCAM | Neuronal Cell Adhesion Molecule | Down |
| CHL1 | Close Homolog of L1 | Down |
| APP | Amyloid precursor protein | Down |
| CHGB | Chromogranin B | Down |
| **iNPH vs PHH** |  |  |
| NELL2 | Neural EGFL-like 2 | Down |
| KLK7 | kallikrein-related peptidase 7 | Down |
| EPHA4 | Ephrin type-A receptor 4 | Down |
| **iNPH vs SAH vs PHH** |  |  |
| CCL4 | C-C motif chemokine ligand 4 | Up |
| **PHH vs SAH** |  |  |
| KLKB1 | plasma prekallikrein | Up |
| SERPIND1 | heparin cofactor II | Up |
| CCL20 | C-C motif chemokine ligand 20 | Up |
| CCL4 | C-C motif chemokine ligand 4 | Up |
| CCL3 | C-C motif chemokine ligand 3 | Up |
| OSM | Oncostatin M | Up |

**Figure 4.**
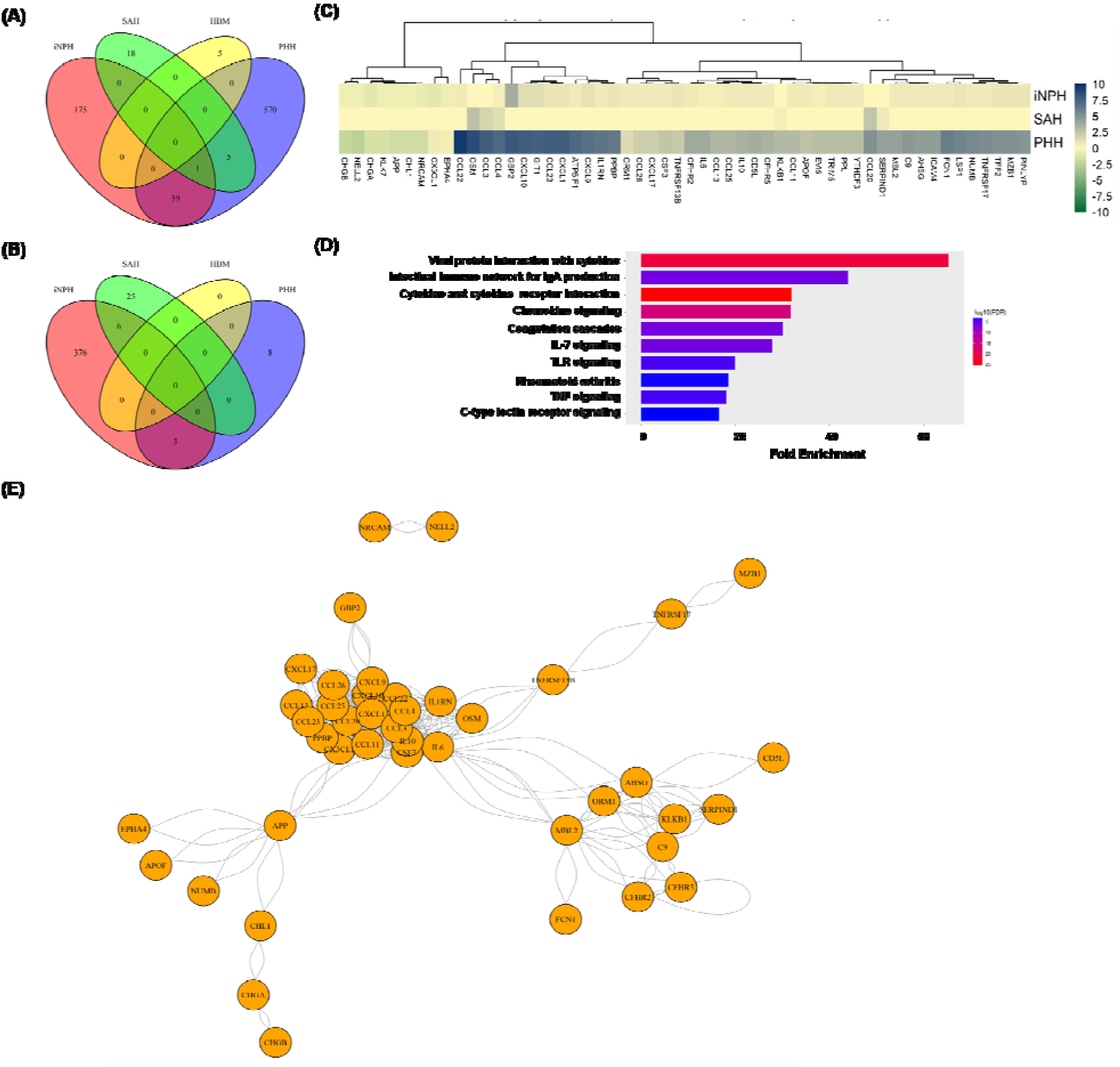
Venn diagrams showing the overlap of (A) upregulated and (B) downregulated proteins among four hydrocephalus subtypes (iNPH, SAH, PHH, and HBM), (C) Heatmap illustrating the expression patterns of the overlapping proteins identified in the Venn analysis across the hydrocephalus subtypes, (D) Top ten enriched pathways for the overlapping protein identified across the hydrocephalus subtypes in KEGG (E) STRING protein-protein interaction network of overlapping proteins

To further delineate the pathways of these overlapping/ commonly associated proteins, we carried out KEGG pathway analysis. The top ten most highly enriched pathways included cytokine– cytokine receptor interaction, IL-17 signaling, TNF signaling, chemokine signaling, coagulation cascades, and TLR signaling, collectively signifying a coordinated activation of innate and adaptive immune networks (Figure 4D). These pathways are well established in the literature as central mediators of neuroinflammatory responses, where elevated cytokine and chemokine production drives leukocyte recruitment and sustained inflammatory signaling in the CNS (e.g., IL-17 and TNF pathways have been implicated in BBB disruption and glial activation **[18, 27]**). Moreover, TLR signaling links pathogen-and damage-associated molecular patterns (PAMPs/DAMPs) to downstream NF-κB activation and proinflammatory cytokine release **[39]**, consistent with pathogen-associated and sterile inflammatory models of hydrocephalus. The enrichment of coagulation cascade pathways further reflects the interplay between inflammation and hemostatic activation, as components of the coagulation system (e.g., thrombin, fibrinogen) which can amplify cytokine production and microglial activation in CNS injury and infection **[14, 56]**. More evidence from STRING protein-protein interaction network confirms the grouped proteins between these signaling cascade of events (Figure 4E). A prominent inflammatory cluster comprised multiple cytokines and chemokines, including IL6, OSM, IL1RN, CCL11, CCL19, CCL20, CCL23, CCL25, CCL26, CXCL11, and CXCL16, indicating extensive connectivity among immune-associated proteins. A separate interconnected module included proteins associated with complement, coagulation, and proteolytic regulation, including C9, KLKB1, SERPIND1, F13B, CTHRC1, and CTHRC-related proteins. APP was positioned within an additional network containing APOF, NUMB, CHI3L1, CHGA, and CHGB, linking proteins associated with neuronal/cellular function and inflammatory processes. Overall, the network exhibited a modular architecture dominated by inflammatory and coagulation-associated proteins, alongside smaller groups of proteins associated with neuronal and cellular processes. Full list of identified DEPs and their protein-protein interaction networks can be found here (https://gowthami-yankan.github.io/protein_network.html). Together, these findings underscore a robust inflammatory milieu involving cytokine networks, PRR (pattern recognition receptor) signaling, and coagulation–immune crosstalk, supporting the concept that dysregulated immune responses contribute to the pathophysiology observed across hydrocephalus subtypes.

### 3.4. Ventricular vs. lumbar CSF proteomic profiles reveal compartment-specific signatures

Due to the limited availability of publicly accessible proteomic datasets derived from vCSF (five studies) and LP CSF (8 studies) across hydrocephalus subtypes (Figure 5A), ventricular and lumbar CSF data within each disease group were integrated to enhance analytical robustness and capture the overall proteomic alterations associated with each hydrocephalus subtypes in this study. However, we performed differential expression analyses relative to appropriately matched control datasets to investigate overlapping and distinct proteomic profiles across different CSF sampling compartments, with emphasis placed on the directionality of protein dysregulation rather than absolute inter-compartmental concentration differences. Given the well-established biological distinctions between ventricular and lumbar CSF **[55]**, potential compartment-specific variability was carefully considered during interpretation to minimize confounding and ensure that conclusions reflected disease-associated rather than sampling-site effects.

**Figure 5.**
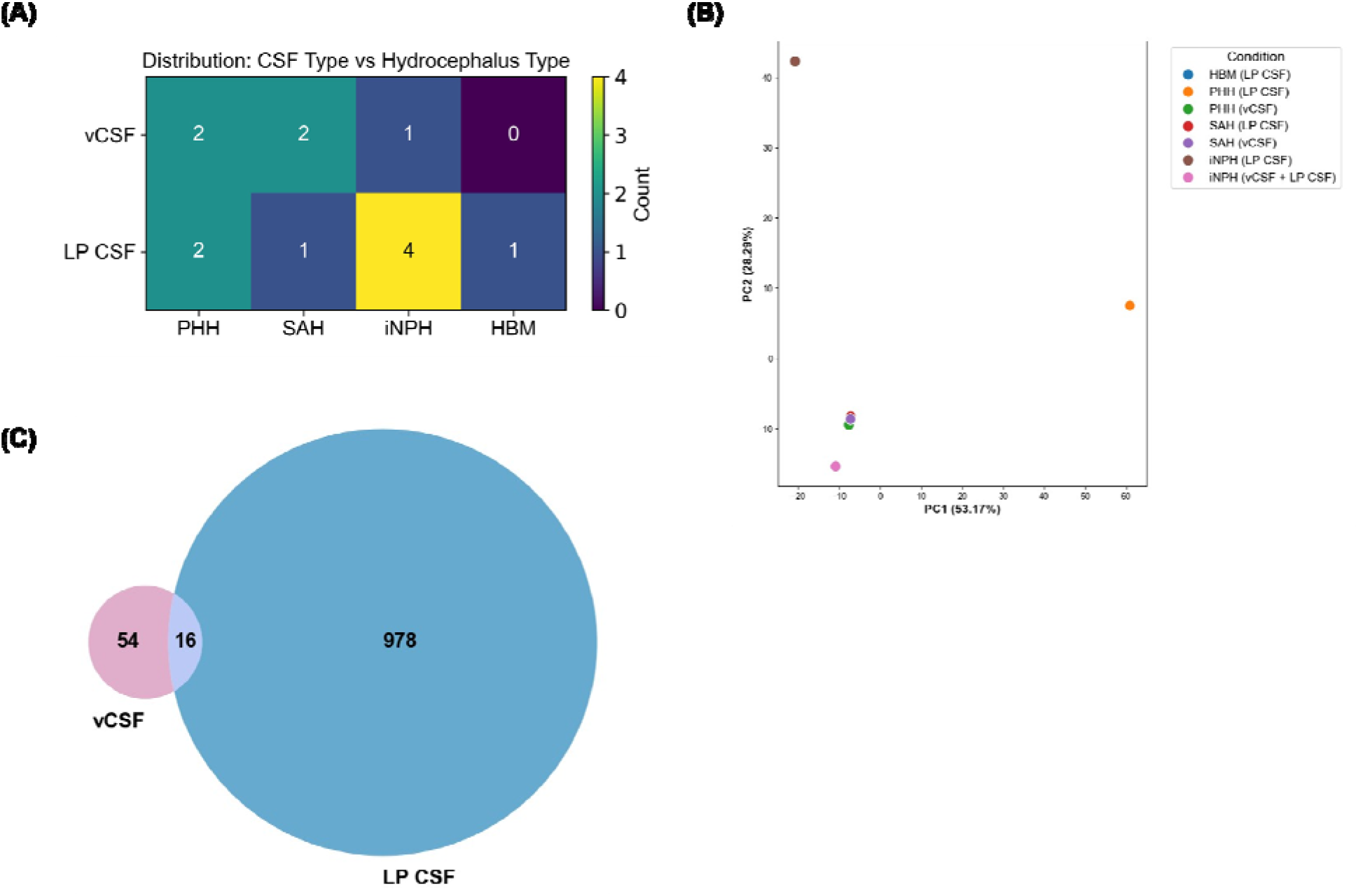
Overlaps between vCSF and LP CSF proteomics study findings. (A) Heatmap showing the number of studies on vCSF and LP CSF in each subtypes of hydrocephalus, (B) Principal component analysis (PCA) of CSF protein profiling across hydrocephalus subtypes. PC1 (53.17%) and PC2 (28.29%) together account for 81.46% of the total variation in the data, (C) Venn diagram illustrating total DEPs from vCSF and LP CSF groups. 16 DEPs were shared between the vCSF and LP CSF profiling

As shown in the PCA plot, (PC1 (53.17%) and PC2 (28.29%)), PHH (LP CSF) and iNPH (LP CSF) groups show distinct separation indicative of the compartment-specific proteomic differences while SAH and PHH ventricular CSF samples cluster more closely, indicative of significant overlap between their proteomic signatures (Figure 5B). Although it would be beneficial to consider LP CSF proteomics of PHH and iNPH groups separately from vCSF proteomics, given the limited number of hydrocephalus proteomic studies currently available for both datasets across multiple hydrocephalus subtypes, independent compartment-specific analyses would substantially reduce statistical and comparative interpretability. While 978 DEPs and 54 DEPs were uniquely identified in LP CSF and vCSF samplings, respectively, 16 DEPs showed shared proteomics profile (Figure 5C), strongly suggesting markedly distinct proteomic profiles between ventricular and LP CSF. Proteins that fell in this overlapping category include ADA, AGRN, APOA4, APOE, APP, CAT, CCL20, CCL3, CCL4, CHL1, CLU, GSN, L1CAM, OSM, PRDX2, SERPINF1 that are broadly associated with neuroinflammation, synaptic organization, neuronal injury, and ECM remodeling, supporting their potential involvement in ventricular pathology and hydrocephalus progression. The substantially lower number of DEPs identified in vCSF should be interpreted cautiously and cannot be attributed solely to biological differences between vCSF and LP CSF. The studies included in this cross-study analysis differed in hydrocephalus etiology, cohort size, proteomic platform, analytical depth, comparison groups, and criteria used to define differential protein abundance, all of which can influence the number of reported DEPs. In addition, fewer vCSF proteomic datasets were available for inclusion than LP CSF datasets, limiting the opportunity to identify recurrent proteins within the ventricular compartment. Nevertheless, the limited overlap between compartments is consistent with previously reported compartment-specific differences in ventricular and LP CSF proteomic composition.

## 4. Discussion

Despite the profound neurological burden, the therapeutic options remain restricted for hydrocephalus patients. Current management still relies on surgical interventions, including CSF shunting and endoscopic procedures, which are frequently accompanied by complications such as infection, obstruction, shunt malfunction, and repeated surgical revisions. While surgery remains the primary treatment modality for hydrocephalus, its limitations have led to the consideration of pharmacological therapies targeting the molecular and cellular mechanisms underlying CSF dysregulation and ventricular enlargement **[6, 26]**. Nevertheless, the development of pharmacological modalities for hydrocephalus has been significantly limited by the lack of understanding of the cellular and molecular mechanisms underpinning CSF secretion and regulation **[6, 40]**. This limitation has driven growing interest in systems-level approaches aimed at characterizing the complex molecular alterations underlying hydrocephalus pathology. In particular, CSF and ChP tissue proteomics have emerged as powerful tools for identifying dysregulated biological pathways, biomarker candidates, and novel therapeutic targets associated with disease progression in recent years **[30, 40, 54]**. Therefore, this manuscript has systematically investigated and compared hydrocephalus proteomics studies (mainly CSF proteomics) published over the past 16 years across different hydrocephalus subtypes (PHH/IVH, SAH, iNPH, HBM) and discussed the candidate biomarkers and the associated signaling pathways linked with disease pathogenesis.

PHH subtype investigated with IVH datasets in preterm infants illustrated high enrichment for blood coagulation cascades, inflammatory signaling involving cytokine, chemokine and IL-17 signaling pathways along with JAK-STAT, PI3K-Akt signaling and MAPK signaling (Figure 3A). The key DEPs linked to these pathways include TGF-β1, MMP-9, TNF-α, IL-6, IL-8, IL-10, CCL-3, APP, p-tau, L1CAM, NELL2, AGRN, FTL, NFL, GFAP, BCAN, and NCAM-1. Similarly, in SAH-associated hydrocephalus mainly from an adult population dataset, significant pathway enrichment was again found in coagulation cascades, TNF signaling, cytokine-chemokine receptor signaling, TLR signaling, leukocyte recruitment, inflammatory signaling and ECM-associated signaling (Figure 3B). The key DEPs linked to these pathways include caspase-I, SAA1, S100B, APLP1, APP, CHGB, THY-1, CCL-3, CCL-4, CCL-20, IL-6, IL-8, IL-10, MCP-1, DNER, OSM, MCP-3 and PTGDS. Further, in iNPH group significantly enriched pathways include cytokine-chemokine receptor signaling, IL-17 signaling, JAK-STAT, PI3K-Akt signaling, ECM-associated signaling (Figure 3C). The key DEPs linked to these pathways include PTPRQ, VIM, L1CAM, PCDH9, NFL, APP, MBP, p-tau, CXCL6, CCL-3, CCL-4, IL-18, IL-8, GFAP, APOE, NRCAM, SCG3 and DAG1. Lastly, HBM group was highly enriched pathways overlapping with PHH/IVH and SAH-associated hydrocephalus groups, including coagulation cascades, IL-17 signaling, proteolysis pathways, and immune-related pathways (Figure 3D). The key DEPs linked to these pathways include AZU1, S100A8, C4BPB, UBE2D1 and TNFRSF13.

As pointed out by most of the proteomic studies across multiple hydrocephalus subtypes, it is evident that there is a consistent increase in cytokines, chemokines, complement proteins, and acute phase-associated proteins involved in inflammatory response in the CSF **[7, 8, 17, 37, 53, 70]**, as indicated in Table A4. This validates that hydrocephalus cannot be viewed solely as a disorder of CSF flow mechanics, but also as a condition in which inflammatory dysregulation within the CSF compartment actively contributes to disease progression. It is important to note that inflammatory processes within the CSF compartment extend into adjacent neural tissue. For example, Yuan et al., showed that elevated IL-1β and TNF-α levels correlate with kallikrein-6 (KLK6) expression, implicating protease mediated inflammatory amplification and synaptic injury **[70]**. This reveals that inflammatory mediators within the CSF not only disrupt fluid homeostasis but may also directly influence neuronal and synaptic integrity and elevate the levels of IL-1β and TNF-α in brain parenchyma. Supplementing this, a recent study from our laboratory showed increased NLRP3 expression in cortical neurons of a genetic rat model of hydrocephalus **[4]**. This evolving body of evidence confirms that hydrocephalus is a disorder that starts with inflammatory CSF homeostasis and extends to the rest of the brain.

**Table A4.** Inflammatory proteins identified in hydrocephalus, measurement methods used and the biological sources.

| Inflammatory Protein | Type of measurement and Source of | Therapeutic | Ref |
| / Molecule | Inflammatory Proteins | Strategy |  |
|---|---|---|---|
| Pro-inflammatory cytokine: IL-1β, TNF-α | ELISA was used to detect protein in the brain parenchyma and CSF of rat communicating hydrocephalus model | Small interfering RNA (siRNA)-mediated knockdown of KLK6 improves synaptic health | [70] |
| CC chemokines: CCL2, CCL5, MIP-1α<br>CXC chemokines: CXCL1, CXCL5<br>Pro-inflammatory Cytokines: TNF-α, IL-1β, IL-6, IFN-γ, IL-17a, IL-18<br>Anti-inflammatory / Regulatory Cytokines: IL-10 | CSF of PIH and PHH rat model using a rat cytokine array / chemokine array 27 plex | Rapamycin (mTOR inhibitor) antagonizes the "cytokine storm" at the choroid plexus and reduces CSF hypersecretion and ventriculomegaly. | [53] |
| CC chemokines: CCL3, CCL4, MCP-3<br>Pro-inflammatory Cytokines: IL-6<br>Anti-inflammatory / Regulatory Cytokines: IL-10 | Markers elevated in CSF samples from human PHH patients were analyzed using the PEA, and choroid plexus tissue from a rat PHH model was analyzed by RNA sequencing (RNA-seq) | The authors suggest that therapeutic targeting of the hemorrhage-induced inflammation detected in both CSF and choroid plexus tissue could be a strategy for future PHH treatments | [37] |
| CC chemokines: MCP-1 and CCL4 | Markers that were increased in the LP CSF of iNPH patients were analyzed using a PEA | None | [7] |
| Pro-inflammatory Cytokines: IL-1β, IL-18<br>Inflammasome effector: Caspase-1 | CSF of Human patients with SAH using ELISA<br>Brain parenchyma of rat model of SAH using western blot and Immunochemistry. | The inhibition of caspase-1 by VX-765 combat neuroinflammation in CSF following SAH | [17] |
| CC Chemokines: CCL3, CCL4, CCL20, CCL22<br>CXC Chemokines: CXCL8 (IL-8), CXCL10, CXCL1<br>Pro-inflammatory Cytokines: IL-6, TNF, IL-1β, IL-18<br>Anti-inflammatory/regulatory cytokines: IL-10<br>Complement Proteins: C3, C5, CFB (Complement Factor B), CFP (Properdin)<br>Acute-Phase Proteins: ORM1, APOA1 | Proteins were detected using high-sensitivity, multi-targeted proteomic profiling of CSF pediatric PHH patients | None | [8] |
| Complement protein 5 | CSF samples of iNPH patients were analyzed using mass spectrometry | None | [22] |

Moreover, an increase in chemokine levels is another crucial recurring feature of the inflammatory CSF milieu in various forms of hydrocephalus, as these are small signaling proteins (a subset of cytokines) that direct the movement of immune cells **[63]**. Increased MCP-1 (CCL2) and CCL4 levels were common to PHH and iNPH, while many other types of chemokines were expressed in PHH and PIH models **[7, 8, 37, 53]**, as indicated in Table A4. These chemokines may contribute to sustained immune cell trafficking at the CSF to brain parenchyma and CSF to the ChP interfaces. In parallel, activation of complement proteins in the CSF could further amplify this innate immune response. As elevated complement components (C3, C5, CFB, and CFP) have been identified in the CSF of pediatric PHH patients **[8]**, this could increase microglial activation and promote membrane attack complex formation, potentially contributing to the progression of hydrocephalus. Complementing these findings, another study identified increased complement component C5 in iNPH using unbiased proteomics, indicating persistent innate immune activation even in a disorder traditionally considered non-inflammatory **[22]**. Together, these studies support the concept that chronic hydrocephalus phenotypes may involve sustained inflammatory activity within the CSF compartment.

From a translational perspective, these inflammatory findings suggest that combining CSF diversion strategies or combination therapies with targeted immunomodulation may improve the outcomes. For example, Robert et al, discovered that mTOR inhibition with rapamycin attenuated cytokine amplification at the ChP and reduced CSF hypersecretion and ventriculomegaly in PIH and PHH rat models **[53]**. Yuan et al, used siRNA-mediated knockdown of KLK6 to reduce cytokine-associated synaptic injury, highlighting the potential of precise immunomodulatory therapeutic approaches targeting specific inflammatory mediators in communicating hydrocephalus **[70]**. Both studies showed that there is hope for therapeutic interventions to combat inflammation during hydrocephalus.

Against this background, our comparative proteomic analysis provides a broader molecular perspective across hydrocephalus etiologies. We identified converging proteomic signatures involving neuroinflammation and immune activation, together with coagulation cascades, ECM remodeling, and neuronal injury pathways across distinct hydrocephalus subtypes. Despite the differences in etiology and patient populations, recurrent pathways involved in cytokine–chemokine signaling, IL-17, JAK-STAT, PI3K-Akt, and coagulation-related pathways across PHH, SAH-associated hydrocephalus, iNPH, and HBM implicates the presence of common pathogenic processes contributing to ventricular pathology and disease progression, while the subtype-specific DEPs indicate that important molecular differences also exist between hydrocephalus conditions, likely reflecting differences in developmental stage, hemorrhagic injury, infection, and chronic neurodegenerative responses. These outcomes further verify the potential utility of CSF proteomics for identifying clinically relevant biomarkers and therapeutic targets in hydrocephalus. However, longitudinal multi-omics studies will be vital for validating these candidate pathways and advancing the development of targeted non-invasive therapies for hydrocephalus.

## 5. Limitations of the study

A major limitation across CSF proteomic studies is the relatively small cohort size, which limits statistical power and the ability to detect robust differences between groups. In addition, many of the biomarkers were evaluated in only one study, which makes it difficult to comment on the external validity of the results. For example, there is no true consensus definition of what constitutes PHH, some studies were not clear as to how they defined PHH beyond ‘dilation of ventricles after IVH. Moreover, the patients in hydrocephalus studies vary significantly due to different age groups of PHH (preterm, neonates, infants, adults), and different CSF sample types (LP CSF vs vCSF) and as such, even studies that looked at the same biomarkers may not be comparable. Moreover, some studies looked prospectively at infants early during IVH and PHH when the ventricles were initially found to be dilated, while some studies looked at infants much later in the course of the disease, making the comparison within the same groups hard. More importantly, considering the distinction between vCSF and LP-CSF samples in proteomics studies could be essential, as these compartments may exhibit markedly different proteomic signatures. However, due to the limited datasets available for vCSF and LP CSF samples, differential expression and pathway enrichment analyses were performed without strict compartment-specific stratification in order to enable broader comparative evaluation in this study. To account for potential compartment-related differences, additional PCA and overlap analyses were conducted to assess the extent of shared and distinct proteomic signatures between vCSF and LP-CSF samples. Yet, it is important to emphasize that the CSF sampling source remains an important consideration when comparing hydrocephalus subtypes and interpreting candidate biomarker profiles.

## Supplementary Materials

Supplementary information is provided as separate Word (.docx) and Excel (.xlsx) files.

## Author Contributions

Conceptualization, G.M., P.K., D.A. and B.B.Y.; Study design, data collection, data analysis and visualization, G.M., and P.K.,; Funding acquisition, B.B.Y.; Writing— original draft, G.M., P.K., D.A., and B.B.Y.; Writing—review and editing, G.M. and B.B.Y. All authors have read and agreed to the published version of the manuscript.

## Institutional Review Board Statement

Not applicable

## Informed Consent Statement

Not applicable

## Data Availability Statement

No new data were generated in this study and this study utilized only previously published proteomics datasets which are cited in the references

## Acknowledgments

Not applicable

## Disclaimer/Publisher’s Note

The statements, opinions and data contained in all publications are solely those of the individual author(s) and contributor(s) and not of MDPI and/or the editor(s). MDPI and/or the editor(s) disclaim responsibility for any injury to people or property resulting from any ideas, methods, instructions or products referred to in the content.

